# How synaptic strength, short-term plasticity, and input synchrony contribute to neuronal spike output

**DOI:** 10.1101/2021.12.21.473708

**Authors:** Alexandra Gastone Guilabert, Moritz O. Buchholz, Benjamin Ehret, Gregor F.P. Schuhknecht

## Abstract

Neurons integrate from thousands of synapses whose strengths span an order of magnitude. Intriguingly, in mouse neocortex, the few ‘strong’ synapses are formed between similarly tuned cells, suggesting they determine neuronal spiking output. This raises the question of how other computational primitives, including ‘background’ activity from the many ‘weak’ synapses, short-term plasticity, and temporal factors contribute to spiking. We combined extracellular stimulation and whole-cell recordings in mouse barrel cortex to map excitatory postsynaptic potential (EPSP) amplitudes and paired-pulse ratios of excitatory synaptic connections converging onto individual layer 2/3 (L2/3) neurons. While net short-term plasticity was weak, connections with EPSPs > 2 mV were exclusively depressing. There was no evidence for clustering of synaptic properties on individual neurons. Instead, EPSPs and paired-pulse ratios of connections converging onto the same cells spanned the full range observed across L2/3, which critically constrains theoretical models of cortical filtering. To investigate how different computational primitives of synaptic information processing interact to shape spiking, we developed a computational model of a pyramidal neuron in the rodent L2/3 circuitry, which was constrained by our own experiments and published *in vivo* data. We found that the ability of strong inputs to evoke spiking depended on their high temporal synchrony and high firing rates observed *in vivo* and on synaptic background activity – and not primarily on synaptic strength, which further amplified information transfer. Our results provide a framework of how cortical neurons exploit complex synergies between temporal coding, synaptic properties, and noise to transform synaptic inputs into output firing.

## Introduction

Pyramidal neurons in neocortex compute spiking responses on the basis of synaptic inputs they receive from thousands of neurons in the surrounding brain tissue. The strengths of these inputs span one order of magnitude and typically follow a lognormal distribution: while the majority of synaptic connections evoke small excitatory postsynaptic potentials (EPSPs), a small minority elicits comparably large EPSPs (Markram et al., 1997; Tarczy-Hornoch et al., 1999; Song et al., 2005; Feldmeyer et al., 2006; Buzsáki and Mizuseki, 2014; Cossell et al., 2015). Intriguingly, in mouse primary visual cortex (V1), such ‘strong’ connections were found to occur predominantly between those neurons that also exhibit the most similar receptive field properties *in vivo* (Cossell et al., 2015). From these observations, a simple organizational principle of synaptic strength was proposed, in which the majority of the synaptic excitation necessary for action potential firing is provided by a small fraction of strong synaptic inputs, which determine the spike output of the postsynaptic neuron (Cossell et al., 2015). The notion that synaptic strength is the primary determinant for the functional properties of neocortical circuits is attractive because it suggests that mapping the strongest connections in functional or structural analyses reveals the true underlying functional organization of neocortical circuits. However, a more complex picture recently emerged from ferret V1, where the response selectivity of neurons to visual stimulation was found to be determined by the cumulative weight of all co-active synapses, and could not simply be predicted from the tuning of synapses with large EPSPs (Scholl et al., 2020).

Several other observations give further weight to the notion that synaptic strength alone is insufficient to explain neuronal response properties. Synapses are complex biophysical devices, whose response during ongoing activation is insufficiently captured by only a single weight parameter. It is intriguing that those cortical synapses that elicit the largest EPSPs also tend to exhibit the most pronounced short-term depression (Reyes and Sakmann, 1999; Jouhanneau et al., 2015; Lefort and Petersen, 2017), which can vastly reduce the total charge a synapse can deliver to its postsynaptic partner during repeated activation (Stratford et al., 1996; Castro-Alamancos and Oldford, 2002; Chung et al., 2002; Abbott and Regehr, 2004; Boudreau and Ferster, 2005; Bruno and Sakmann, 2006). Thus, synaptic connections with large EPSPs recorded *in vitro* may operate in a significantly depressed state *in vivo* due to ongoing spontaneous and stimulus-evoked activation (Boudreau and Ferster, 2005). Furthermore, even the largest EPSP amplitudes provide only a fraction of the depolarizing charge necessary to drive the membrane potential of a cortical neuron through the spike threshold. Thus, temporal coincidence in presynaptic spike trains must necessarily be an important factor for information coding in neocortex (Bruno and Sakmann, 2006; Banitt et al., 2007; Wang et al., 2010; Schoonover et al., 2014; Scholl et al., 2020). Finally, neurons *in vivo* operate in the presence of significant synaptic background activity. Spontaneous firing rates of pyramidal cells in the superficial layers of rodent sensory areas range between 0.08 to 0.39 Hz *in vivo* (Waters and Helmchen, 2006; de Kock et al., 2007; Kerr et al., 2007; de Kock and Sakmann, 2009; Niell and Stryker, 2010, 2010; O’Connor et al., 2010). Because pyramidal neurons in rodent sensory areas are estimated to receive input from up to ~8000 synapses (Schüz and Palm, 1989), they must experience hundreds to thousands of spontaneous synaptic events per second. In rodent V1, synaptic connections with small EPSPs occur predominantly between cells that display different response properties and thus fire with little temporal synchrony during visual stimulation (Cossell et al., 2015). Thus, in rodent sensory areas, the vast majority of excitatory synapses formed with any given pyramidal neuron provide a constant bombardment of excitation that seems relatively unrelated to the tuning of that neuron. Therefore, to compute spiking responses from their synaptic inputs, neocortical neurons operate in a complex parameter space. While much research has been conducted on the computational role of synaptic strength [e.g. (Lefort et al., 2009; Cossell et al., 2015; Scholl et al., 2020)], short-term plasticity [e.g. (Abbott et al., 1997; Castro-Alamancos and Oldford, 2002; Chung et al., 2002; Banitt et al., 2007; Rothman et al., 2009; Díaz-Quesada et al., 2014)], and the temporal structure within synaptic inputs [e.g. (Bruno and Sakmann, 2006; Banitt et al., 2007; Wang et al., 2010; Schoonover et al., 2014)], it remains much less studied how these parameters act together to shape information transfer in sensory areas.

Here, we combined experimental work and data-driven computational modeling to investigate systematically how this complex parameter-space could shape the spiking responses of pyramidal neurons in L2/3 of mouse barrel cortex (S1). The distributions and patterns of action potential firing rates (de Kock et al., 2007; de Kock and Sakmann, 2009; Sakata and Harris, 2009; O’Connor et al., 2010), synaptic strength (Lefort et al., 2009; Cossell et al., 2015; Seeman et al., 2018), correlations within neuronal activity (Kerr et al., 2007; Sato et al., 2007), and temporal correlations within synaptic inputs converging onto the same neuron (Cossell et al., 2015) have been well-characterized for L2/3 in rodent sensory areas *in vivo*. However, even though paired-pulse ratios have been measured for excitatory synapses across all cortical layers and different areas and species, most studies relied on small datasets that aimed to detect general differences in the mean (Reyes and Sakmann, 1999; Feldmeyer et al., 2006; Costa et al., 2013; Jouhanneau et al., 2015; Lefort and Petersen, 2017; Seeman et al., 2018). Thus, a detailed characterization of the exact statistical distribution of short-term plasticity in mouse sensory L2/3 is missing. Likewise, the relationship between synaptic strength and short-term plasticity has not been characterized clearly for L2/3. Finally, it remains unknown whether synaptic connections that converge onto the same neuron exhibit a systematic bias of EPSP amplitudes (Koulakov et al., 2009) or short-term plasticity, which could endow individual neurons with low-pass filter or high-pass filter properties, if they were to receive predominantly depressing or facilitating synapses, respectively (Chance et al., 1998; Fortune and Rose, 2000, 2001; Abbott and Regehr, 2004). We addressed these questions by combining whole-cell recordings of L2/3 pyramidal neurons in barrel cortex slices with extracellular stimulation of putatively single axons of passage. Then, we developed a computational model of a L2/3 pyramidal neuron that received excitatory inputs from 270 other L2/3 neurons (Sarid et al., 2013), whose synaptic strengths and short-term plasticity were modeled after our experimental data. Presynaptic inputs were set to display temporal firing patterns constrained by *in vivo* data: the few synaptic connections eliciting large EPSPs fired temporally correlated spikes at high frequencies and were termed ‘strong’ inputs, while the more numerous connections triggering small EPSPs – termed ‘weak’ inputs – fired uncorrelated spikes at lower frequencies (Cossell et al., 2015). By selectively manipulating the relationship between synaptic strength, short-term plasticity, and temporal structure in the synaptic inputs, we characterized the importance of each of these parameters and their interdependencies in our simulation.

## Results

### Mapping synaptic strength and short-term plasticity in L2/3

We characterized the distribution of EPSP amplitudes and corresponding paired-pulse ratios of excitatory synaptic connections formed with regular-spiking neurons in barrel cortex L2/3 and tested the theoretical prediction that synaptic connections converging on the same postsynaptic cell may have systematically biased strengths (Koulakov et al., 2009) or short-term plasticity properties. In order to be able to characterize multiple, different synaptic connections formed with a given L2/3 neuron, we measured somatic whole-cell responses to extracellular paired-pulse stimulation of single axons at multiple locations in the surrounding L2/3 (Fig. 1 A, B).

**Figure 1.**
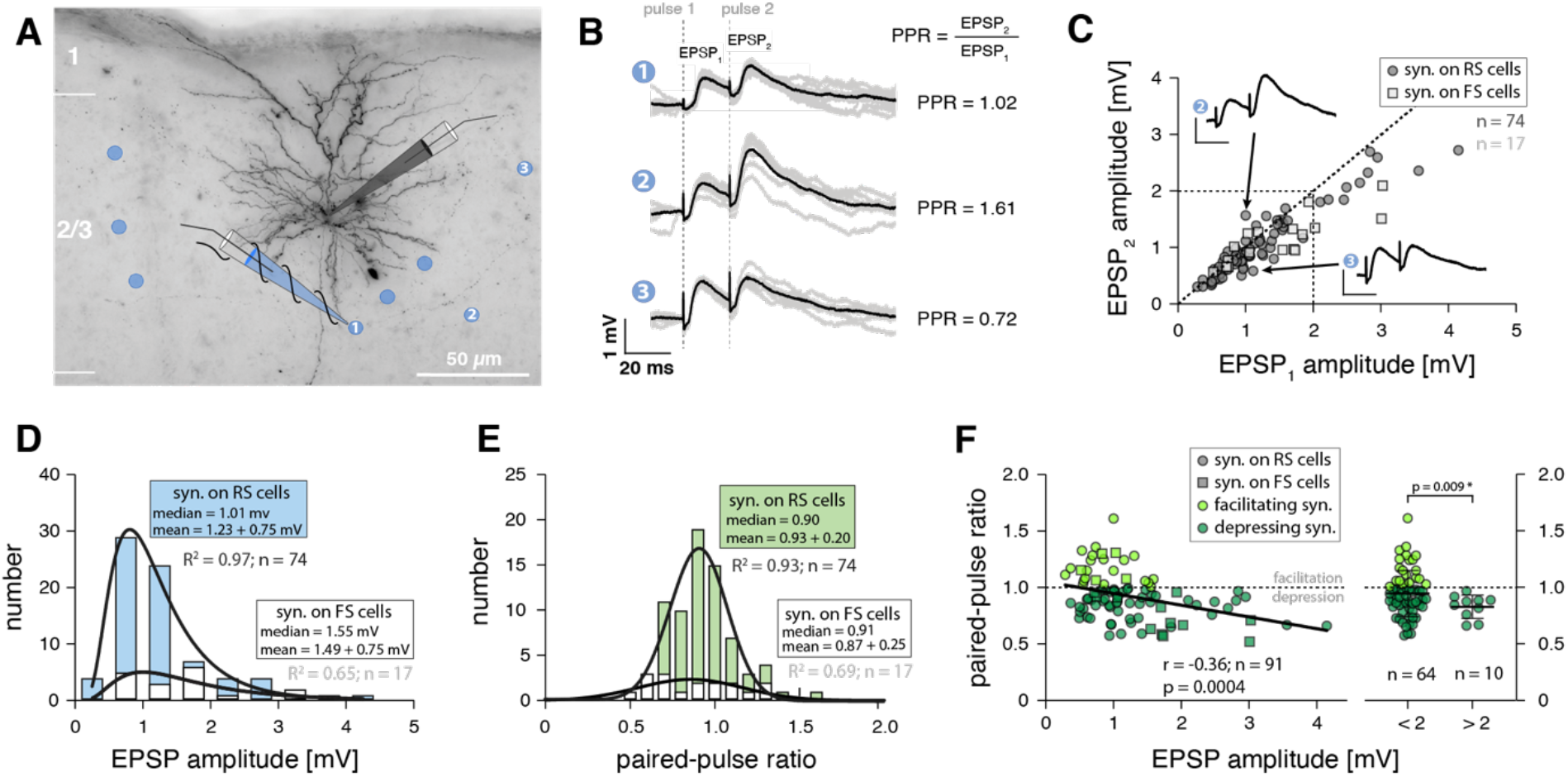
EPSP amplitudes and paired-pulse ratios of excitatory synaptic connections in barrel cortex L2/3. **A** Example of recorded regular-spiking L2/3 neuron in mouse barrel cortex visualized through post-hoc biocytin histology. Blue dots indicate locations of successful extracellular stimulation, blue pipette signifies extracellular stimulation electrode. The neuron’s responses to stimulation at three different positions (labeled 1-3) are shown in B. **B** Somatic voltage recordings following 20 ms paired-pulse stimulation in the locations indicated by numbers. Grey traces, individual trials; black traces, average response; paired-pulse ratios (PPR) indicated. For timing of extracellular stimulation pulses (dashed lines), note the electrical stimulation artifact in somatic voltage responses. **C** Scatter plot showing, for all recorded excitatory synaptic connections, the responses to the second pulse versus the response to the first pulse (corresponding to the EPSP amplitude) of the paired-pulse stimulation paradigm. Circles, synaptic connections formed with regular-spiking (RS) neurons; squares, connections formed with fast-spiking neurons (FS). Data points below diagonal indicate depressing synaptic connections, dots above diagonal indicate facilitating connections. Voltage traces, same as traces 2 and 3 in B with identical same scale bars. **D** Distribution of EPSP amplitudes recorded in regular-spiking (blue) and fast-spiking L2/3 (white) L2/3 neurons; histograms were fit with lognormal functions (R^2^, goodness of fit). **E** Distribution of paired-pulse ratios recorded in regular-spiking (green) and fast-spiking L2/3 (white) L2/3 neurons; histograms were fit with Gaussian functions (R^2^, goodness of fit). **F** Left, scatter plot showing relationship of EPSP amplitude and paired-pulse ratio for excitatory synaptic connections formed with regular-spiking (circles, n = 74) and fast-spiking (squares, n = 17) cells; light green, facilitating connections; dark green, depressing connections. Non-parametric Spearman correlation statistics indicated; line was fit to all datapoints with linear regression. Right, comparison of paired-pulse ratios of synaptic connections formed with regular-spiking neurons that were binned into ‘small’ (EPSP < 2 mV) and ‘large’ (EPSP > 2 mV) synaptic connections (parametric Welch’s t test).

We obtained recordings from 20 regular-spiking neurons for which we identified a total of 74 sites at which minimal extracellular stimulation evoked EPSPs (mean of 3.7 synaptic connections per neuron). For a subset of these regular-spiking cells, we performed post-hoc biocytin histology to confirm that they were indeed pyramidal neurons (Fig. 1 A; see *Methods*). Additionally, we recorded from 4 fast-spiking neurons (i.e., interneurons, as confirmed by post-hoc histology) for which we found a total of 17 extracellular stimulation sites (4.3 synaptic connections per neuron). Thus, our complete dataset contained 91 evoked EPSPs recorded across 24 L2/3 neurons. We applied stringent quality controls to ensure that we activated single axons of passage with our minimal stimulation protocol (see *Methods*) and that we did not stimulate the same axon of passage multiple times. Briefly, we only included synaptic connections for which the smallest observable EPSP was evoked in an all-or-none manner in a fraction of trials and if the mean EPSP amplitude and failure rate remained constant throughout the recording (Larkman et al., 1991; Allen and Stevens, 1994). Moreover, different synaptic connections converging onto the same postsynaptic cell were only included when their location of stimulation was > 50 µm away from all previous stimulation locations.

The distribution of peak amplitudes across the 74 EPSPs recorded in regular-spiking cells ranged from 0.29 mV to 4.15 mV (mean ± s.d.: 1.23 ± 0.75 mV), was markedly right-skewed, and could be fit well with a lognormal distribution (R^2^ = 0.97) (Fig. 1 D). The mean coefficient of variation was 0.19 ± 0.06, the mean EPSP onset latency was 2.14 ± 1.12 ms and the mean 10 – 90% rise time was 2.54 ± 0.86 ms. For all 74 synaptic connections, we also recorded the paired-pulse ratio at an inter-spike interval of 20 ms. Interestingly, the distribution of paired-pulse ratios appeared noticeably symmetrical with a mean ± s.d. of 0.93 ± 0.20 and could be fit well with a normal distribution (R^2^ = 0.93) (Fig. 1 E).

A similar picture emerged for the 17 EPSPs recorded in the fast-spiking cells: their amplitudes ranged from 0.52 mV to 3.03 mV (mean ± s.d.: 1.49 ± 0.75 mV) and were best captured by a lognormal distribution (R^2^ = 0.65) (Fig. 1 D). The mean coefficient of variation was 0.18 ± 0.06, the mean onset latency was 2.44 ± 0.99 ms, and the 10 – 90% rise time was 0.83 ± 0.4 ms. The distribution of corresponding paired-pulse ratios was also markedly symmetrical with a mean of 0.87 ± 0.25 (Fig. 1 E) and could be fit well with a normal distribution (R^2^ = 0.69).

Given their different symmetries (lognormal versus normal, respectively), the question arose of how EPSP amplitudes and their corresponding paired-pulse ratios could be mapped onto one another, i.e., whether there was a systematic relationship between synaptic strength and short-term plasticity. Interestingly, a scatter plot of the response amplitudes to the 2^nd^ stimulation pulse against the response amplitudes to the 1^st^ stimulation pulse (corresponding to the EPSP amplitude) showed the tendency that synaptic connections with larger EPSPs were depressing, while connections with smaller EPSPs exhibited a range of facilitating and depressing paired-pulse ratios (Fig. 1 C, F). However, there was no significant correlation in our dataset between EPSP amplitude and short-term plasticity for connections formed with either regular-spiking or fast-spiking neurons. A significant negative correlation only emerged when we pooled all synaptic connections recorded in the study (Fig 1 F). Thus, EPSP amplitude and short-term plasticity appeared to be only weakly correlated across a larger number of excitatory synaptic connections in L2/3. Because of the limited number of synaptic connections recorded in fast-spiking neurons, however, we excluded these data from further analysis and focused the rest of our study on the synaptic connections recorded in regular-spiking neurons.

To investigate this question further, we binned our dataset of synaptic connections recorded in regular-spiking neurons depending on their EPSP amplitude (into 0.5 mV bins, not shown). Critically, we found that in all bins with EPSP amplitudes below 2 mV, synaptic connections displayed a range of facilitating and depressing paired-pulse ratios (not shown). By contrast, all connections with EPSP amplitudes above 2 mV were depressing (n = 10) (Fig. 1 F). When we split the dataset accordingly, we found that connections below 2 mV had a mean paired-pulse ratio of 0.95 ± 0.20 (i.e., exhibiting little net short-term plasticity), while connections above 2 mV had a significantly lower mean paired-pulse ratio of 0.83 ± 0.10 (Fig. 1 F).

### No clustering of connections with similar paired-pulse ratios on L2/3 neurons

Next, we investigated whether EPSP amplitudes and short-term plasticity across those synaptic connections formed with the same regular-spiking L2/3 neurons followed the same distributions as those of all 74 connections across all regular-spiking neurons. Alternatively, the synaptic inputs onto a given cortical neuron may be statistically correlated, i.e. individual neurons could receive synaptic connections with systematically biased EPSP amplitudes or paired-pulse ratios that deviate from the overall distributions found across L2/3, which may constitute a mechanism to endow individual cells with high-pass or low-pass filtering properties (Fortune and Rose, 2001; Abbott and Regehr, 2004). For a total of 8 regular-spiking neurons, we were able to characterize at least 5 different afferent synaptic connections (47 connections in total, mean of 5.9 connections per cell). We will refer to the distribution of paired-pulse ratios and EPSP amplitudes across all our recorded synapses as the “population distribution” and to the distributions of paired-pulse ratios and EPSPs of synaptic connections converging onto a single cell as “cell distributions”. We used the non-parametric Kolmogorov-Smirnov test to detect if there was a significant difference between the respective cell distributions and the population distribution. Interestingly, for all 8 cells, the cell distributions were not significantly different from the population distribution for both EPSP amplitude and paired-pulse ratios (Fig. 2 A, B).

**Figure 2.**
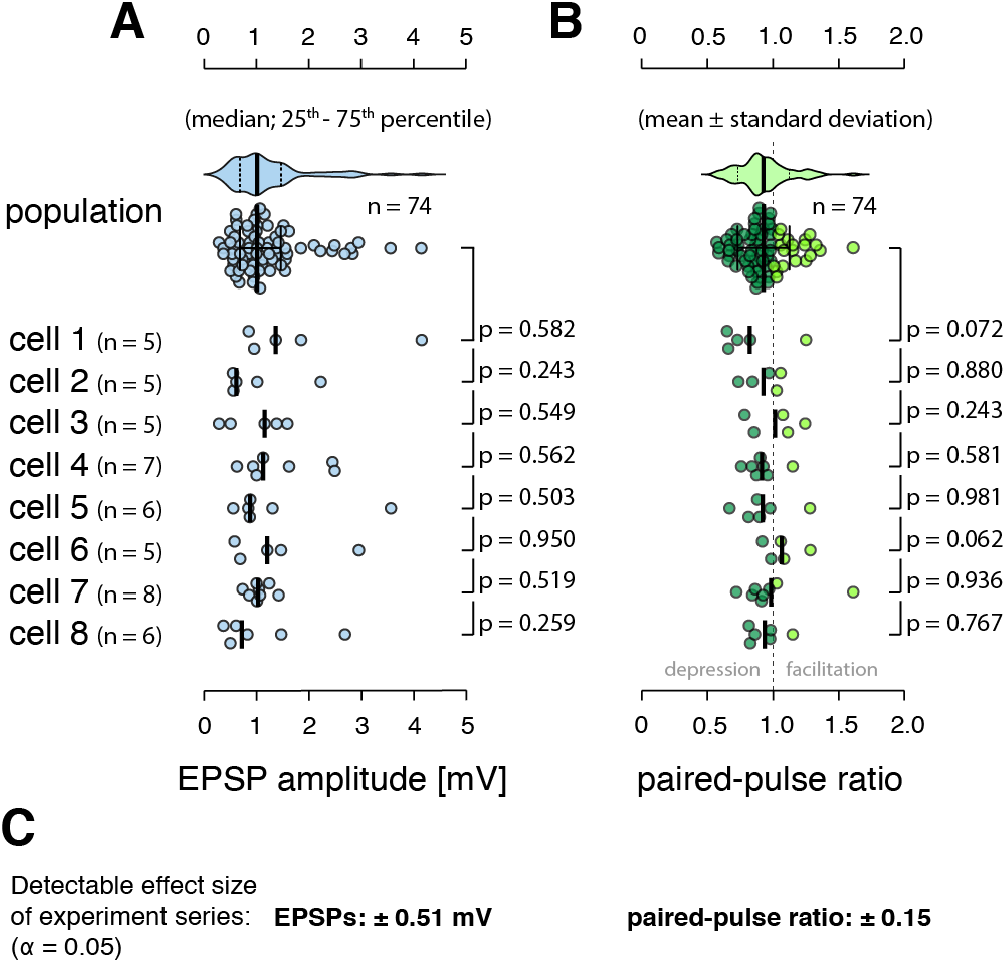
Excitatory synaptic connections formed with regular-spiking L2/3 neurons do not exhibit a systematic clustering of EPSP amplitude and short-term plasticity. **A** Top, distribution of EPSP amplitudes recorded across all regular-spiking neurons (population distribution). Bottom, distributions of the EPSP amplitudes across the 8 regular spiking neurons, for which at least 5 synapses were found (cell distributions). N, number of synapses recorded per cell; p, non-parametric Kolmogorov-Smirnov test between each cell distribution and the population distribution, medians are indicated. **B** Representation of short-term plasticity data, panel layout as in A; light green, facilitating synaptic connections; dark green, depressing connections, means are indicated. **C** Estimation of the effect sizes that are detectible across the experimental series at a 5% significance level.

Precise quantification of synaptic short-term plasticity requires electrophysiological recordings. Using whole-cell patch-clamp recordings in combination with minimal stimulation of axons of passage, however, limits the number of synaptic connections that can be recorded for any given neuron, yielding low statistical power on the level of individual cells. Therefore, we conducted a power analysis to estimate the detectable effect sizes in our dataset (see *Methods* for details). For detecting a significant (α = 0.05) difference between each of the 8 paired-pulse ratio cell distributions and the population distribution, the Kolmogorov-Smirnov test had an average power of 17% for an effect size of 0.1, a power of 53% for an effect size of 0.2, and a power of 85% for an effect size of 0.3, where effect size corresponds to a systematic difference in the means of the cell distributions. Thus, the statistical power was low on the level of individual experiments. Because we could repeat the experiment 8 times, however, even small systematic differences between cell distributions and population distribution, while undetectable in single experiments, should have been revealed in at least one or a few of the 8 neurons we recorded from. To investigate this further, we used a binomial model (see *Methods*) to assess the power of the entire experimental series by asking: what systematic difference in paired-pulse ratios should have been observed in at least one of the 8 experiments at the 95% significance levelã We found that the probability to detect a significant difference across our entire dataset was 78% for an effect size of 0.1 and 99.7 % for an effect size of 0.2, with the 95% significance level at an effect size of 0.15. Critically, an effect size of 0.15 is below the paired-pulse ratio difference of 0.16 that we detected between the small- and large-EPSP connections formed with pyramidal neurons in L2/3 (Fig. 2 C). Thus, our experimental series achieved the statistical power necessary to detect differences in paired-pulse ratios at physiological magnitudes that we found to exist in L2/3. This suggests that short-term plasticity of excitatory synapses formed with individual regular-spiking cortical neurons in L2/3 spans the full range observed in L2/3 and is not markedly functionally clustered on the level of single neurons.

Likewise, for detecting a significant difference between each of the 8 EPSP cell distributions and the population distribution, the Kolmogorov-Smirnov test had an average power of 4.9% for an effect size of 0.2 mV, a power of 15% for an effect size of 0.4 mV, and a power of 46% for an effect size of 0.6 mV. Analogous Monte Carlos simulations showed that the probability of detecting a significant difference in the mean EPSP amplitudes across our entire dataset was 72% for a systematic effect size of 0.4 mV and 99.3 % for a systematic effect size of 0.6 mV, with the 95% significance level at 0.52 mV (Fig. 2 C). In summary, these are important experimental results that contradict the theory-inspired hypothesis that synaptic inputs onto single cortical neurons may be statistically correlated (Koulakov et al., 2009).

### Modeling the interplay of synaptic strength, short-term plasticity, and temporal input structure

We generated a two-compartment, conductance-based model of a L2/3 pyramidal neuron to investigate how synaptic strength, short-term plasticity, and temporal structure in synaptic inputs interact within the L2/3 circuitry to shape the response properties of cortical neurons (Fig. 3 A – C; *see Methods for details)*. For this purpose, we developed a data-driven modeling approach: we constrained firing rates and pairwise correlations of presynaptic inputs by *in vivo* observations and synaptic strength and short-term plasticity by our experimental data recorded in regular-spiking neurons.

**Figure 3.**
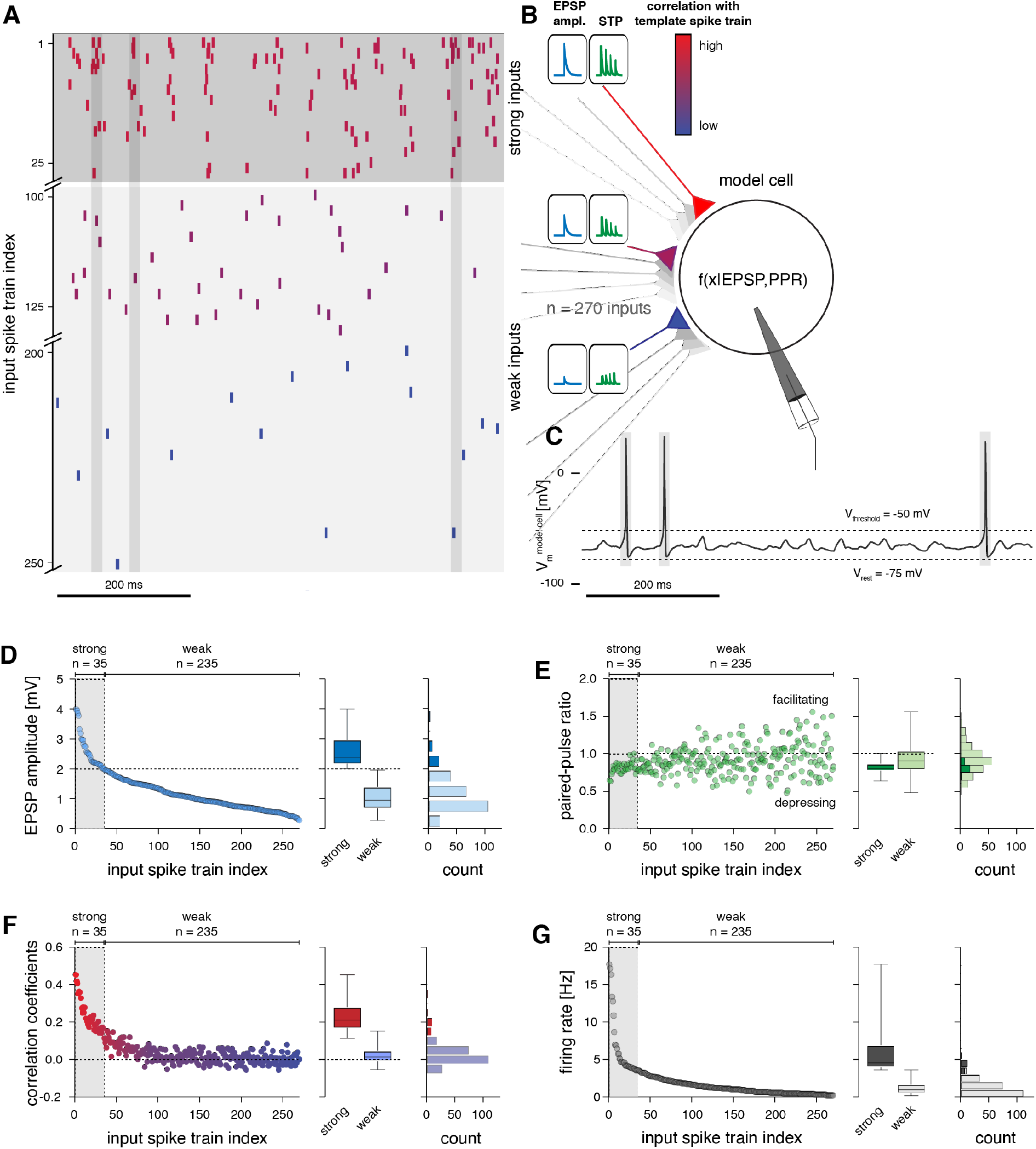
Default setup of the L2/3 neuron model. **A** Example of input spike trains fed to the model cell. Strong inputs (top) fired with higher frequencies and temporal correlation (color coded), compared to weak inputs (bottom). Vertical grey bands indicate the resulting spike timing in the model cell (same as in C). Note that some weak inputs did not spike in the depicted 200 ms time window because of their low firing rates. **B** Strong inputs were set to have larger EPSP amplitudes and corresponding short-term depression, while weak inputs were set to evoke smaller EPSPs and correspondingly weak net short-term plasticity, in accordance with our *in vitro* recordings. **C** Simulated membrane potential of model neuron following activation with the input spike trains shown in A. **D** Left, EPSP amplitudes across the 270 input spike trains. Center, comparison of EPSP amplitudes between strong and weak inputs (median, 25 – 75 % percentile, and ranges are indicated). Right, same data plotted as histogram. **E** Left, 20 ms paired-pulse ratios across the 270 input spike trains. Center, comparison of paired-pulse ratios between strong and weak inputs (median, 25 – 75 % percentile, and ranges are indicated). Right, same data plotted as histogram. **F** Left, Pearson correlation coefficients of the 270 input spike trains with the template spike train that was used to generate the pairwise correlation structure (see *Methods*); color code as in A, B. Center, comparison of correlation with template spike train between strong and weak inputs (median, 25 – 75 % percentile, and ranges are indicated). Right, same data plotted as histogram. **G** Left, firing rates of the 270 input spike trains. Center, comparison of firing rates between strong and weak inputs (median, 25 – 75 % percentile, and ranges are indicated). Right, same data plotted as histogram. The relationships between parameter distributions reflects our *in vitro* data and *in vivo* data adopted from Cossell et al. (2015).

Briefly, the model neuron received excitatory inputs from 270 presynaptic neurons (Feldmeyer et al., 2006; Sarid et al., 2013), whose synaptic weights (Fig. 3 D) and short-term plasticity properties (Fig. 3 E) were constrained following our extracellular stimulation experiments (see *Methods*). Note that this number of presynaptic L2/3 cells is based on the assumption that L2/3 neurons form on average 3 anatomical synapses with their postsynaptic partners in L2/3 (Feldmeyer et al., 2006; Sarid et al., 2013). In our model, this is captured by the fact that the axons of passage we activated with minimal stimulation must have also formed multiple synapses with the recorded neurons on average. This is evident when comparing the range of EPSP amplitudes we recorded with minimal stimulation (0.29 – 4.15 mV) with EPSP amplitudes obtained from paired recordings (0.15 – 2.25 mV), for which the number of anatomical synapses per connection (mean of 1.6) was additionally established from EM (Holler et al., 2021). The temporal input correlations (Cossell et al., 2015) (Fig. 3 F) and the firing rates (O’Connor et al., 2010) (Fig. 3 G) across the 270 synaptic inputs were constrained by published *in vivo* data for rodent cortex (see *Methods*), such that a small number of strong synaptic inputs fired temporally correlated spikes at high frequencies and exhibited large EPSP amplitudes and corresponding short-term depression. The remaining majority of weak synapses, providing ‘background’ activity, were set to fire at low frequencies and in a temporally uncorrelated pattern, resembling a random Poisson process, and exhibited low EPSP amplitudes without pronounced net short-term plasticity (Fig. 3 A, B).

After the model was set up in this manner, we verified that all parameters were distributed following experimental data and that the interdependencies between EPSP amplitude and short-term plasticity and EPSP amplitude and temporal correlation structure (Cossell et al., 2015) were preserved (Fig. 4). The model EPSP amplitude distribution (Fig. 3 D, Fig. 4 A; mean ± s.d.: 1.23 ± 0.69 mV, n = 270) and paired-pulse ratio distribution for a 20 ms paired-pulse interval (Fig. 3 E, Fig. 4 B; mean ± s.d.: 0.91 ± 0.19, n = 270) did not differ from the distributions we had measured in regular-spiking neurons *in vitro* (p = 0.90 and p = 0.67, respectively; non-parametric Kolmogorov-Smirnov tests). The mapping between EPSP amplitude and short-term plasticity across the model inputs (Fig. 4 A, B) followed the same relationship as observed *in vitro*: EPSP amplitudes > 2 mV had significantly lower paired-pulse ratios (mean ± s.d.: 0.82 ± 0.08) compared with EPSP amplitudes < 2 mV (Fig 3 E; mean ± s.d.: 0.92 ± 0.20; p < 0.0001, parametric Welch’s t test), and due to the larger sample size compared to our *in vitro* data, there was a negative correlation between EPSP amplitude and paired-pulse ratio (r = -0.26, p < 0.0001, n = 270, non-parametric Spearman correlation coefficient). In accordance with electrophysiological recordings obtained from rodent sensory L2/3 *in vivo* (O’Connor et al., 2010), the firing rates of the inputs followed a lognormal distribution (R^2^ = 0.88) with a mean of 1.9 ± 2.4 Hz (Fig. 4 C), the strong synaptic inputs had a mean firing rate of 6.4 ± 4.1 Hz (maximum: 17.7 Hz), and the weak synaptic inputs had a mean firing rate of 1.2 ± 0.9 Hz (Fig. 3 G). The strong inputs exhibited the highest pairwise correlation coefficients (mean ± s.d.: 0.24 ± 0.09; range: 0.11 to 0.45), while the weak inputs exhibited little correlation (mean ± s.d.: 0.02 ± 0.04; range: -0.05 to 0.15)(Fig. 3 F)(Cossell et al., 2015).

**Figure 4.**
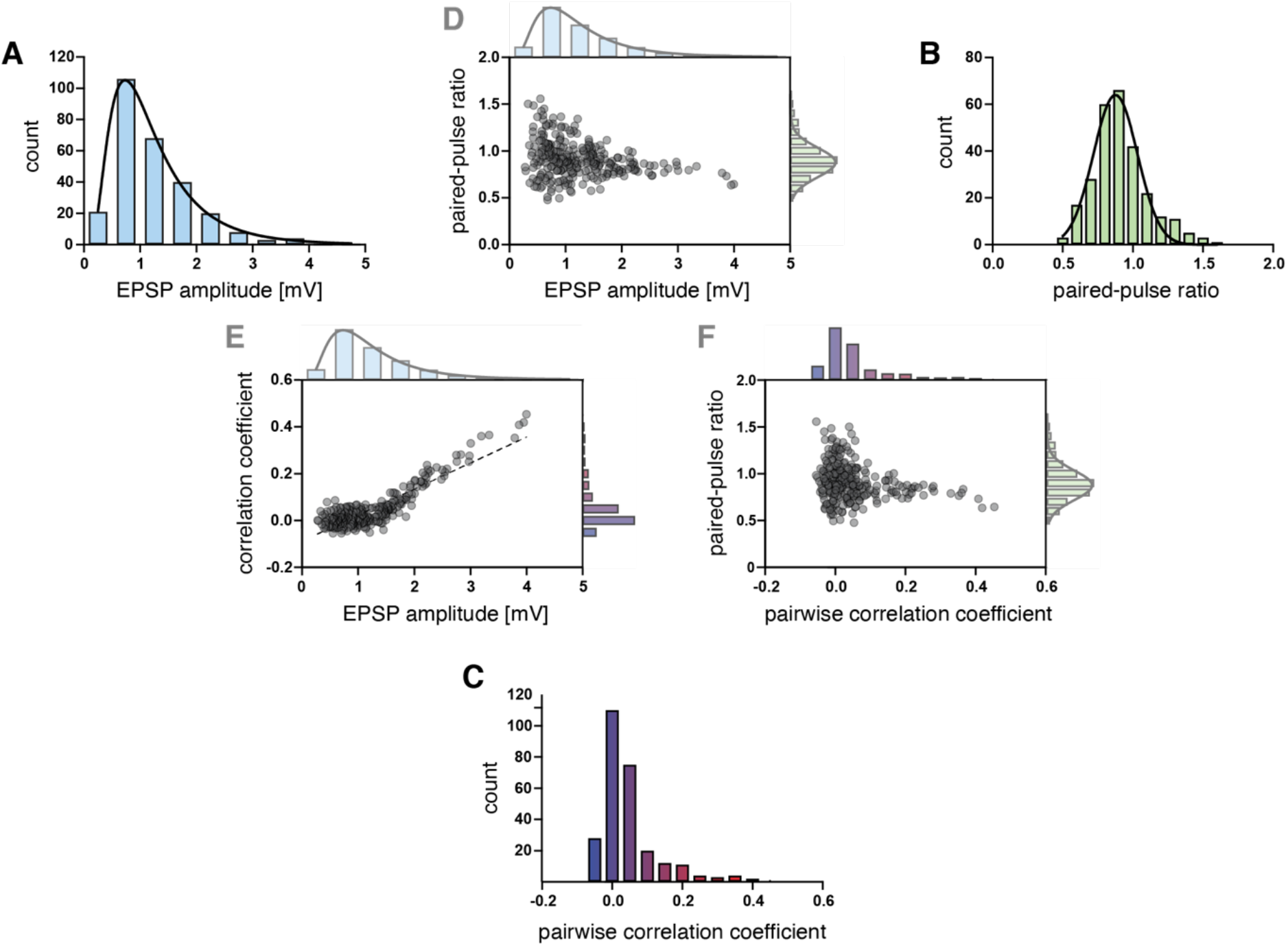
Mapping between synaptic strength, short-term plasticity, and correlation in input spike trains. **A** EPSP distribution for 270 inputs generated from our *in vitro* recordings. **B** 20 ms paired-pulse ratio distribution for 270 inputs generated from our *in vitro* recordings. **C** Pairwise-correlation coefficients for 270 inputs generated from *in vivo* data adopted from Cossell et al. (2015). **D** Scatter plot of relationship between EPSP amplitudes and 20 ms paired-pulse ratios for the 270 inputs. **E** Scatter plot of relationship between EPSP amplitudes and pairwise-correlation coefficients for the 270 inputs. **F** Scatter plot of relationship between 20 ms paired-pulse ratios and pairwise-correlation coefficients for the 270 inputs.

To examine information transfer between the synaptic inputs and the output firing pattern of the model neuron, we measured the Pearson correlation coefficient between each input spike train and the model neuron’s output spike train. We further characterized the neuronal gain of the model cell by mapping its input-output relationship (i.e., the probability of spiking as a function of the number of coincident synaptic inputs). By selectively manipulating the relationship between synaptic strength, short-term plasticity, and temporal structure in the synaptic inputs, we then systematically characterized the contribution of each of these parameters on information transfer and neuronal gain. Each experiment was repeated for a total of 100 simulation runs; whereby for each iteration, we randomly re-generated a new set of 270 input spike trains.

First, we ran the simulation in its default ‘physiological’ setup, i.e., with parameters and parameter-mappings as found in our *in vitro* recordings and published *in vivo* data (Fig. 3, 5). Critically, without further tuning, the model neuron reproduced key properties of rodent L2/3 pyramidal neurons *in vivo*. It generated output spike trains with an average firing rate of 4.81 ± 0.71 Hz (Fig. 3 C, 5 F), which is in excellent agreement with experimental measurements of *in vivo* spike rates in mouse barrel cortex L2/3 (O’Connor et al., 2010). The average membrane voltage (V_m_) of the model neuron was -65.93 mV ± 7.82 mV (Fig. 5 B, D), comparable to *in vivo* whole-cell recordings in mouse L2 (Jouhanneau et al., 2015). As expected, the strong synaptic inputs shared the highest Pearson correlation coefficients (mean ± s.d.: 0.11 ± 0.038; range: 0.064 to 0.20) with the resulting output spike train of the model neuron (Cossell et al., 2015), while the weak inputs displayed correlation coefficients one order of magnitude smaller (mean ± s.d.: 0.012 ± 0.013; range: -0.0041 to 0.065) (Fig. 5 E). Across all inputs, spike trains with decreasing intrinsic correlation, smaller EPSP amplitudes, and lower spike rates displayed increasingly lower correlation coefficients with the output spike train (Fig. 5 E). We confirmed that the Pearson correlation coefficients indeed detected correlations in spike timing rather than in firing rates by randomizing the output spike times following a random Poisson process while keeping the output firing rate identical. Reassuringly, the correlations between all inputs and the output spike train then dropped to -0.0005 ± 0.0032 (not shown).

**Figure 5.**
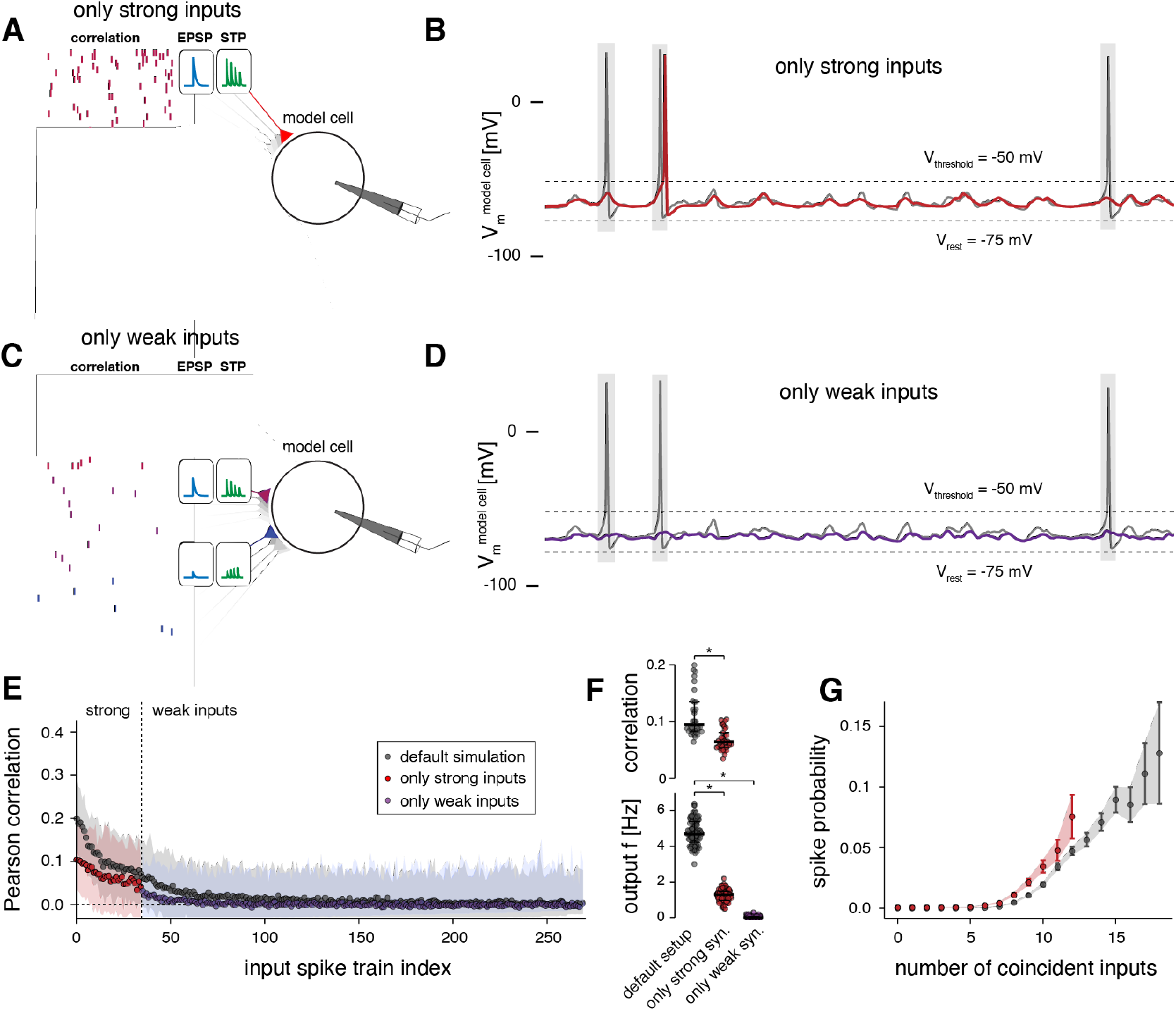
Uncorrelated activity from weak inputs enhances information transfer of strong synaptic inputs. **A** Schematic of model setup with weak inputs removed. **B** Example spike train of the model cell in its default setup (grey) and when weak inputs are removed (red). **C** Schematic of model setup with strong inputs removed. **D** Example spike train of the model cell in its default setup (grey) and when strong inputs are removed (purple). **E** Pearson correlation coefficients of the 270 input spike trains with the output spike train of the model cell. Results of three model setups are shown: default simulation (all inputs, as in Fig. 3) and setups introduced in A, C. Shaded regions, 95 % confidence bounds for correlation coefficients obtained from 100 runs of the simulation. **F** Top, Pearson correlation coefficients between the strong synaptic inputs and the output spike train of the model neuron for the default simulation and setup introduced in A. Bottom, output firing rate of model cell for the default simulation and the setups introduced in A-C. (Data are averages across 100 simulation runs; median and 25 – 75 % percentile indicated; non-parametric Kolmogorov-Smirnov test, * p < 0.0001.) **G** Probability of output spiking as a function of coincident spikes across all input spike trains. (Note that the maximum number of coincident inputs within the 20 ms measurement window was 12 when only strong inputs were included, thus determining the maximum x-value for the red curve.)

### Synaptic background activity enhances information transfer of strong inputs

We probed the relative influence of the strong versus weak synaptic inputs on the output spiking of our model cell. Critically, when we removed the weak inputs (Fig. 5 A, B), the mean correlation between the strong inputs and the output spike train was reduced to 0.068 ± 0.019 (range: 0.034 to 0.10) (Fig. 5 E, F). The output firing rate of the model neuron dropped to 1.26 ± 0.34 Hz (Fig. 5 F) and its average V_m_ was hyperpolarized to -68.39mV ± 4.57mV. Despite the sharp drop in information transfer of the strong synaptic inputs, those inputs with the highest intrinsic correlation and synaptic strength still maintained the highest correlation with output spiking (Fig. 5 E). Removal of weak inputs also resulted in a steeper slope of the input-output curve (Fig. 5 G), confirming that synaptic ‘background noise’ has a divisive effect on neuronal gain. This noise broadens a neuron’s sensitivity to the range of temporal correlations in input spike trains by increasing the time window over which coincident inputs can be integrated to evoke spiking, a finding in agreement with previous studies (Silver, 2010).

Conversely, when we removed the strong synaptic inputs from the simulation (Fig. 5 C), uncorrelated activity provided by the weak inputs was by itself unable to drive the postsynaptic neuron above spiking threshold and the output firing rate dropped to 0.045 ± 0.068 Hz (Fig. 5 F). This is because the 235 weak inputs fired at an average frequency of 1.2 ± 0.9 Hz with mean EPSP amplitudes of 1.03 ± 0.42 mV, which resulted in a mean membrane potential of 67.68 ± 1.67 mV that rarely crossed the spike threshold (Fig. 5 D). Thus, uncorrelated activity of weak synapses alone was incapable of evoking spikes and did not transfer information encoded in its own spike trains (Fig. 5 E). Importantly, however, it had a powerful computational effect on neuronal activity because it enhanced information transfer of the strong, correlated inputs by a factor of 2.

### Output spiking requires correlation and high firing rates of strong inputs

Next, we decoupled the high temporal correlation and high firing rates of strong inputs from their larger synaptic strengths by randomly assigning the EPSP amplitudes and their corresponding short-term plasticity properties across the input spike trains (Fig. 6 A). Note that the original coupling between EPSP amplitude and short-term plasticity was maintained in this experiment, i.e., synapses with larger EPSPs still exhibited depression and synapses with smaller EPSPs exhibited facilitation. When the model was set up in this manner, the firing rate of the output neuron decreased to 1.11 ± 0.68 Hz (Fig. 6 B, D). Critically, inputs with higher temporal correlation and higher firing rates still contributed more strongly to the firing of the model neuron (Pearson correlation mean ± s.d.: 0.056 ± 0.016; range: 0.035 to 0.095) compared to inputs with lower temporal correlations and lower firing rates (mean ± s.d.: 0.009 ± 0.008; range: -0.004 to 0.042) (Fig. 6 C). This means that synaptic strength by itself did not determine which inputs transmitted the most information to the spike train of the output neuron. Instead, in our simulation, the combination of high temporal correlation and elevated firing rates of strong synaptic inputs was the primary determinant for evoking correlated spiking in the output neuron. However, matching larger EPSP amplitudes to inputs that fired with high temporal correlation and high firing rates (i.e., our default setup), as observed for the strong synaptic inputs *in vivo* (Cossell et al., 2015), increased their correlation with the spike train of the model neuron by a factor of 2 and enhanced their information transfer (Fig. 6 C, D). Decoupling the large EPSP amplitudes from the correlated inputs (by shuffling EPSP amplitudes amongst all input spike trains) furthermore resulted in a flatter slope of the model’s input-output curve and a reduced responsiveness to coincident inputs (maximum spike probability (P_max_) of 0.15; Fig 6 E). This suggests that assigning the largest EPSP amplitudes to those inputs that fired at high temporal correlation has a multiplicative effect on neuronal gain, leading to signal amplification as a mechanism to increase efficient information transmission of strong inputs (Silver, 2010).

**Figure 6.**
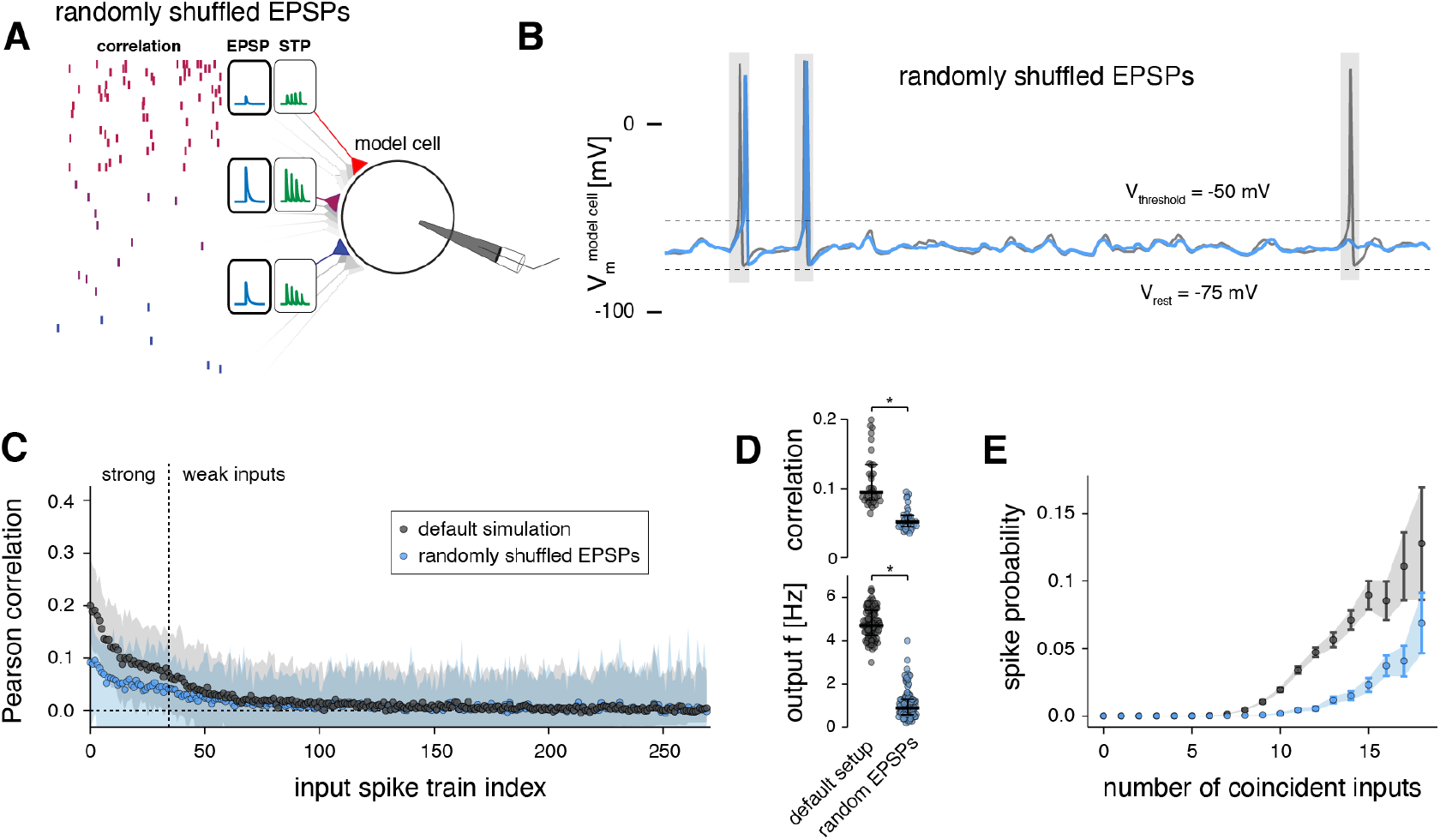
Temporal correlation and firing rates primarily determine output spiking, synapse strength enhances response. **A** Schematic of model setup with shuffled EPSP amplitudes; note that the relationship of EPSP amplitude and short-term plasticity was maintained. **B** Example spike train of the model cell in its default setup (grey) and with shuffled EPSP amplitudes (blue). **C** Pearson correlation coefficients of the 270 input spike trains with the output spike train of the model cell. Results of two model setups are shown: default simulation (as in Fig. 3) and setup introduced in A. Shaded regions, 95 % confidence bounds for correlation coefficients obtained from 100 runs of the simulation. **D** Top, Pearson correlation coefficients between the strong synaptic inputs and the output spike train of the model neuron for the default simulation and setup introduced in A. Bottom, output firing rate of model cell for the default simulation and the setup introduced in A. (Data are averages across 100 simulation runs; median and 25 – 75 % percentile indicated; non-parametric Kolmogorov-Smirnov test, * p < 0.0001.) **E** Probability of output spiking as a function of coincident spikes across all input spike trains.

### Short-term plasticity balances the computational effects of strong and weak inputs

Next, we removed the short-term plasticity mechanism from all synapses, such that they exhibited paired-pulse ratios of 1 for all inter-spike-interval durations, i.e., synaptic strength remained static during repeated stimulation (Fig. 7 A, B).

**Figure 7.**
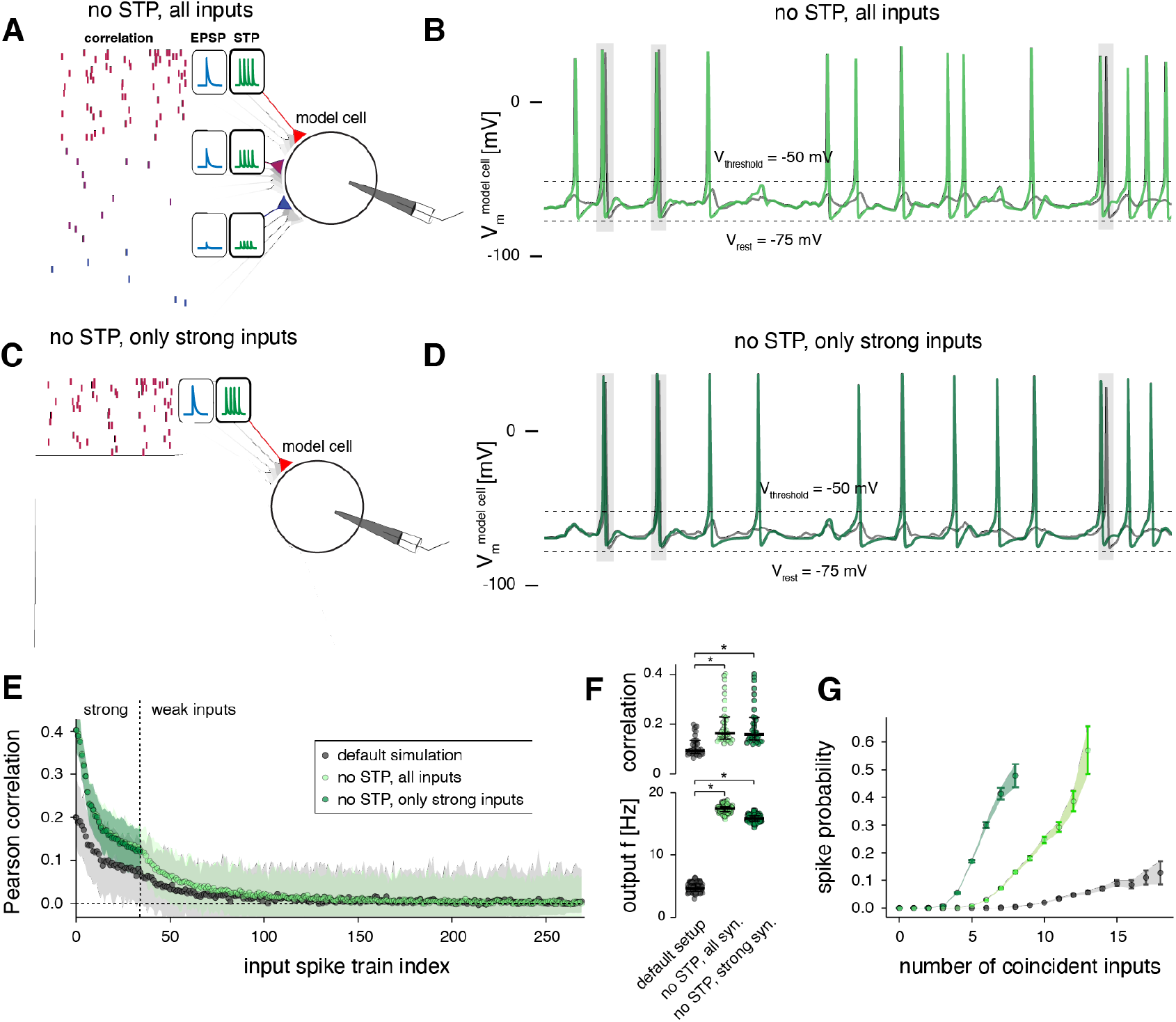
Short-term plasticity balances the computational effects of strong and weak inputs. **A** Schematic of model setup with short-term plasticity mechanisms removed; note that all spike trains exhibit a paired-pulse ratio of 1. **B** Example spike train of the model cell in its default setup (grey) and when short-term plasticity mechanisms are removed (light green). **C** Schematic of model setup with short-term plasticity mechanisms removed and the weak inputs removed in addition. **D** Example spike train of the model cell in its default setup (grey) and when short-term plasticity mechanisms and weak inputs are removed (dark green). **E** Pearson correlation coefficients of the 270 input spike trains with the output spike train of the model cell. Results of three model setups are shown: default simulation (as in Fig. 3) and setups introduced in A, C. Shaded regions, 95 % confidence bounds for correlation coefficients obtained from 100 runs of the simulation. **F** Top, Pearson correlation coefficients between the strong synaptic inputs and the output spike train of the model neuron for the default simulation and setups introduced in A-C. Bottom, output firing rate of model cell for the default simulation and the setups introduced in A-C. (Data are averages across 100 simulation runs; median and 25 – 75 % percentile indicated; non-parametric Kolmogorov-Smirnov test, * p < 0.0001.) **G** Probability of output spiking as a function of coincident spikes across all input spike trains.

When running the simulation in this setup, the model neuron fired at 17.4 ± 0.63 Hz, which was an even higher frequency than exhibited by those input spike trains with the highest firing rates (Fig. 7 F). At the same time, the mean correlation coefficient of the strong inputs with the output spike train of the model neuron had doubled to 0.20 ± 0.082, with the largest values exceeding 0.4 (Fig. 7 E, F). Also the correlation coefficients of the weak inputs with the output spike train had increased to 0.018 ± 0.023 (Fig. 7 E). Notably, because the weak synaptic inputs had been only mildly depressing on average in our default setup (Fig. 3 E), removing their short-term plasticity mechanism should only have a small net boosting effect on their total excitatory drive. To confirm this, we additionally removed the weak inputs from the model entirely (Fig. 7 C, D) and found that this indeed had no significant effect on the correlation coefficients between the strong inputs and the resulting spike train of the model neuron (Fig. 7 E, F) nor on the output firing rate of the model cell (Fig. 7 F). Thus, after removing short-term plasticity, the computational effect of the weak inputs in maximizing information transfer of strong inputs had become entirely redundant. In this regime, the strong synapses alone could determine the spiking properties of the model neuron.

Furthermore, the slopes of the input-output curves were markedly steeper when the short-term plasticity mechanism was removed and when the weak inputs were removed in addition (Fig. 7 G), which confirms that short-term depression of strong inputs has a divisive impact on neuronal gain (Abbott et al., 1997; Rothman et al., 2009), therefore broadening the neuron’s responsiveness to temporal correlations in input spike trains (Silver, 2010).

## Discussion

We combined experimental work and computational modeling to investigate how the spiking responses of L2/3 pyramidal neurons are shaped by the complex parameter-space of temporal structure within synaptic inputs, synaptic strength, and short-term plasticity.

As a first step, we mapped experimentally the distribution of synaptic strength and short-term plasticity in barrel cortex L2/3. We found that short-term plasticity follows a symmetrical distribution with large variance and is mildly depressing on average. Interestingly, synaptic strength and short-term plasticity were only weakly negatively correlated across our dataset. Instead, their relationship was well-captured by the simple rule that synaptic connections with EPSP amplitudes below 2 mV span the full range of depression and facilitation and exhibit no pronounced average short-term plasticity. By contrast, connections with EPSP amplitudes above 2 mV are exclusively depressing, which raises the intriguing question of what the computational role is for depression of strong synapses.

Our computational model of a L2/3 neuron suggested that the ability of the strong synaptic inputs to evoke spiking in postsynaptic cells relies predominantly on their high temporal correlation and high firing rates, as well as on synaptic background activity from the numerous weak synapses, and not primarily on their synaptic strength. Pairing those ‘driving’, co-tuned synaptic connections with strong synaptic weights, however, as has been reported for rodent V1 (Cossell et al., 2015), does amplify their ability to transmit information to the output spike train.

### Technical considerations of minimal stimulation

We used minimal stimulation of axons of passage to map EPSP amplitudes and paired-pulse ratios of multiple, different synaptic connections formed with the same postsynaptic neurons. This study design rendered paired whole-cell recordings unfeasible, as the number of synaptic connections that can be identified with paired recordings is usually low. To ensure that EPSPs originated from single axons, we carefully followed established protocols (Larkman et al., 1991; Allen and Stevens, 1994) and imposed strict data inclusion criteria (see *Methods*). Minimal extracellular stimulation of axons of passage is a classical technique in neuroscience [e.g. (Larkman et al., 1991, 1992; Allen and Stevens, 1994; Volgushev et al., 1995; Stratford et al., 1996)] and although it cannot be ruled out that several afferent axons may be activated in principle, previous work suggests that this is unlikely to occur in practice. For example, studies that mapped synaptic connections with minimal stimulation have reported similar EPSP amplitudes [(Stratford et al., 1996; Hardingham and Fox, 2006); our data] and numbers of release sites (Hardingham and Fox, 2006) when compared with paired recordings of the same connections (Stratford et al., 1996; Silver et al., 2003; Hardingham et al., 2010; Holler et al., 2021). Thus, the possibility of synaptic connections arising from several afferent axons was assumed to be negligible in our dataset.

### Short-term plasticity in barrel cortex L2/3

While mild average depression in rodent barrel cortex L2/3 is in agreement with previous reports (Reyes and Sakmann, 1999; Feldmeyer et al., 2006), other studies have found excitatory L2/3 synapses in sensory areas to be moderately facilitating on average (Jouhanneau et al., 2015; Lefort and Petersen, 2017; Seeman et al., 2018). Intriguingly, Jouhanneau et al. (2015) and Lefort and Petersen (2017) used paired recordings in layer 2 (L2). While Seeman et al. (2018) conducted paired recordings across the entire thickness of L2/3, they reported only mild average facilitation with large overall heterogeneity.

We recorded from neuronal somata in superficial L2/3, likely corresponding to the layer investigated by Jouhanneau et al. (2015) and Lefort and Petersen (2017), but stimulated axons of passage across the entire depth of L2/3. Thus, differences between these datasets may indicate differences in synaptic properties between L2 recurrent connections (Jouhanneau et al., 2015; Lefort and Petersen, 2017) and the L3 -> L2 pathway. This is in line with a growing body of literature describing structural (Karimi et al., 2020) and functional (Crochet et al., 2011; Petersen and Crochet, 2013) differences between the neuronal circuits in L2 and L3 and further supports the notion that L2 and L3, which are routinely considered to constitute a single computational entity, may in fact possess different computational properties (Petersen and Crochet, 2013; Karimi et al., 2020).

### No evidence for a statistical bias of synaptic innervation on L2/3 neurons

Interestingly, we found no statistical bias of synaptic strength or short-term plasticity of synaptic connections formed with the same pyramidal neurons in L2/3. Instead, our data suggest that synaptic inputs formed with a given L2/3 neuron are not markedly correlated, but that their strengths and short-term plasticity instead follow the same distribution as that of all synaptic connections across the neuropil. Such statistical biases have been hypothesized to explain lognormal firing rate distributions in cortex (Koulakov et al., 2009) and have been proposed as a potential mechanism for endowing neurons with high-pass or low-pass filter properties that may underlie integration and differential activation (Lisman, 1997; Fortune and Rose, 2001; Abbott and Regehr, 2004). Importantly, by demonstrating the absence of such systematic biases on the single-cell level, our experimental results provide a critical biological constraint for theoretical models of how these particular computations may arise in L2/3.

Because we have characterized synaptic strength and short-term plasticity through somatic whole-cell recordings, we cannot exclude the intriguing possibility that statistical biases of synaptic innervation may exist on the level of dendritic branches, in which case such computations may be implemented on a sub-cellular level (Kastellakis et al., 2015; Bloss et al., 2018). Further experiments will be necessary to investigate this possibility.

### L2/3 neuron model reproduces key computation properties of cortical circuits

To address the computational role of depression of strong connections and to investigate how synaptic strength, short-term plasticity, and temporal properties in presynaptic spike trains within the L2/3 circuitry shape the firing properties of neurons, we generated a simplified model of a L2/3 pyramidal neuron and systematically manipulated these parameters in our simulation. The synaptic inputs to the model neuron were constrained by physiological data obtained from our own *in vitro* recordings and with *in vivo* data adopted from the literature (O’Connor et al., 2010; Cossell et al., 2015). To focus our study on the L2/3 circuitry, we constrained the model neuron to receive synaptic connections only from other L2/3 neurons, i.e., connections from layer 4 and the deep layers were not modeled, such that it was not necessary to include inhibitory synapses in the simulation to balance excitation. Reassuringly, without further parameter tuning, the model exhibited key computational properties of cortical neurons that have been characterized in experiments and simulations before: the model cell produced sparse firing at around 5 Hz, which is in excellent agreement with the average spike rate reported for mouse barrel cortex L2/3 (O’Connor et al., 2010), and its output spike train exhibited the highest temporal correlation with the strong synaptic inputs (Cossell et al., 2015). In addition, our simulations could reproduce the effects of multiplicative gain modulation through synaptic background activity (Salinas and Sejnowski, 2001; Chance et al., 2002) and through short-term depression of strong synapses (Abbott et al., 1997; Rothman et al., 2009).

We found that synaptic background activity carried through the weak synapses contributed critically to information transfer of strong inputs through a stochastic resonance-type effect (Faisal et al., 2008): while being incapable of evoking spiking by itself, the weak inputs enabled the model neuron to operate in a regime in which the cell became sensitive and responsive to coincident strong inputs (Bulsara et al., 1991; Hô and Destexhe, 2000; Chapeau-Blondeau and Rousseau, 2002; London et al., 2002; McDonnell and Abbott, 2009; Durand et al., 2013). Even then, the high firing rates and the synchronous activity of multiple strong synapses were needed to evoke spiking in the model neuron (Bruno and Sakmann, 2006; Banitt et al., 2007; Wang et al., 2010; Schoonover et al., 2014; Martin and Schröder, 2016). Notably, synaptic strength alone did not determine which presynaptic cells could evoke spikes (Scholl et al., 2020).

### Short-term depression balances the synaptic drive of strong inputs

The computational role of the relationship between short-term plasticity and synaptic strength has not been addressed in detail in studies of cortical processing. Interestingly, the pronounced short-term depression we observed for synaptic connections eliciting large EPSPs *in vitro* proved necessary to counterbalance the high firing rates, high temporal correlations, and large EPSP amplitudes of strong inputs during ongoing stimulation and was critical for maintaining the responsiveness of the postsynaptic neuron towards input spike trains with the highest temporal correlation. This suggests that short-term depression could act as one of the mechanisms that prevent runaway excitation in the recurrent L2/3 circuitry.

### A framework for orientation tuning in columnar and ‘salt-and-pepper’ cortices

The notion that the minority of strong synaptic inputs determines the response properties of cortical neurons (Cossell et al., 2015; Znamenskiy et al., 2018; Goetz et al., 2021) has recently been challenged by apparently conflicting findings made in V1 of the ferret (Scholl et al., 2020). In mouse V1, neurons with the most similar receptive field properties *in vivo* also formed the strongest synaptic connections with each other, as assessed *in vitro* (Cossell et al., 2015). By contrast, the response selectivity of neurons in ferret V1 *in vivo* was shown to be determined by the cumulative weight of all driving synapses – weak and strong. Intriguingly, the response selectivity could not be predicted from the tuning of strong synapses alone (Scholl et al., 2020).

Our result that spiking in the model neuron was driven predominantly by high temporal input correlation and high firing rates, while synaptic strength further enhanced information transfer of these driving inputs may provide a framework to reconcile these apparently contradictory findings. In the columnar V1 of carnivores (Hubel and Wiesel, 1962), presentation of simple visual stimuli activates populations of neighboring neurons within the same orientation column (Ohki et al., 2005). The axons of pyramidal cells in the superficial layers of V1 form a primary cluster of synaptic boutons around their own somata (Martin et al., 2014). Thus, unlike in rodents, these neurons are excited by many neighboring neurons with the same orientation tuning and ocular dominance. Therefore, ‘columnar’ orientation maps, which are found in visual areas of higher mammals (Gilbert and Wiesel, 1989; Malach et al., 1994; Bosking et al., 1997; Sincich and Blasdel, 2001) may provide the basis for the “strength by numbers” necessary to generate tuned responses (Scholl et al., 2020), without the additional requirement of stronger synapses between co-tuned neurons. Our finding that the high temporal correlation and firing rates of strong inputs, and *not* their larger synaptic strength primarily drive spiking supports this idea and is consistent with the observation that spikes in cat V1 are phase-locked with the local field potential, which reflects synchrony within local neuronal populations (Martin and Schröder, 2016).

By contrast, the ‘salt-and-pepper’ organization of rodent V1 (Girman et al., 1999), means that oriented stimuli activate a spatially diffuse network (Ohki et al., 2005). Therefore, neurons may receive fewer synaptic connections overall from similarly tuned cells and temporal correlation and firing rates alone may be insufficient to achieve orientation tuning. Our observation that pairing large EPSP amplitudes with correlated input spike trains further enhances the capacity of driving inputs to transmit information suggests that this predicted ‘lack of strength by numbers’ in rodent V1 may be compensated for by stronger synapses between similarly tuned neurons (Cossell et al., 2015). This, however, leads to the prediction that in mouse V1, the temporal structure in input spike trains from similarly tuned neurons also plays a key role in generating orientation tuning *in vivo*, a prediction that could be tested experimentally.

In summary, our results provide a framework for how cortical neurons could utilize interactions between the biophysical properties of chemical synapses, the temporal structure of input spike trains, and ‘noise’ in neuronal networks for efficient computation.

## Methods

### Animals

Cortical slices were obtained from 13 male B6/C57 mice between 22 and 29 postnatal days of age under the license of Kevan A.C. Martin (Institute of Neuroinformatics, University of Zurich & ETH Zurich, Zurich, Switzerland). Animal handling and experimental protocols were approved by the Cantonal Veterinary Office, Zurich, Switzerland.

### Slice preparation

Animals were anesthetized with isoflurane, decapitated, and their brains were removed quickly and immersed in ice-cold slicing artificial cerebrospinal fluid (ACSF, containing, in mM: 87 NaCl, 75 sucrose, 26 NaHCO_3_, 10 glucose, 7 MgSO_4_, 2.5 KCl, 1 NaH_2_PO_4_, and 0.5 CaCl_2_, continuously oxygenated with 95% O_2_, 5% CO_2_). Coronal slices containing the barrel cortex were cut at a thickness of 300 μm on a vibratome and transferred to a chamber containing recoding ACSF (containing, in mM: 119 NaCl, 26 NaHCO_3_, 10 glucose, 1.3 MgSO_4_, 2.5 KCl, 1.25 NaH_2_PO_4_, and 2.5 CaCl_2_, continuously oxygenated with 95% O_2_, 5% CO_2_). The slices were kept in recording ACSF at room temperate until the recordings.

### Electrophysiology

Patch pipettes (pipette resistance: 5-7 MΩ, pipette tip diameter: 2 μm) were pulled from borosilicate glass using a P-97 puller (Sutter Instruments) and filled with intracellular solution (containing in mM: 105 K-gluconate, 20 KCl, 10 Na-phosphocreatine, 2 Mg-ATP, 2 Na-ATP, 0.3 GTP, and 10 HEPES, pH was set to 7.2 with KOH). Biocytin (0.5%) was added to the intracellular solution to stain the recorded neurons. Whole-cell patch-clamp recordings were obtained at 34-36 °C from visually identified L2/3 neurons in barrel cortex under an Olympus BX61W1 microscope equipped with infrared differential-interference contrast optics and a 10x and a 60x water-immersion objective. Data were acquired with a Multiclamp 700A amplifier (Axon Instruments), sampled at 10 kHz, filtered at 3 kHz (Digidata 1322A, Axon Instruments) and monitored with the software pClamp (Molecular Devices). We did not add GABA^A^ (Allen and Stevens, 1994; Volgushev et al., 1995; Hardingham and Fox, 2006) or NMDA antagonists to the bath (Allen and Stevens, 1994; Volgushev et al., 1995), as previous studies have not reported any discernible effects on the EPSP waveform from including these blockers in minimal extracellular stimulation experiments (Larkman et al., 1991, 1992, 1997).

Following break-in, the access resistance was typically in the range of 15-30 MΩ and recordings with an access resistance > 30 MΩ were discarded. The bridge potential was compensated and liquid-junction potential was not corrected. V_m_ after break-in ranged from -85 to -70 mV. If V_m_ drifted during recordings, a holding current was injected to keep the membrane at its initial resting potential, which was rarely necessary. Because V_m_ was close to the reversal potential of GABA^A^ in all experiments, we expect there was no contamination of our recorded EPSPs by inhibitory connections.

We then performed minimal stimulation of single axons of passage according to established protocols (Larkman et al., 1991; Allen and Stevens, 1994), as follows. After establishing whole-cell recordings, we identified presynaptic axons forming synapses with the recorded cells by carefully moving a monopolar extracellular stimulation electrode (filled with ACSF) through L2/3 at an oblique angle and delivering repeated 0.1 ms current pulses of 10-12 μA amplitude using an A360 stimulator (World Precision Instruments) until an EPSP was detected in the patched neuron. Synaptic connections were typically detected when the stimulation electrode was located 20-400 μm distant from the soma of the recorded cell. To achieve stimulation of single axon fibers synapsing onto the patched neuron, we then decreased the stimulation amplitude until the EPSP was not elicited anymore and subsequently increased the stimulation amplitude until the smallest observable EPSP was evoked reliably in an all-or-none manner in a fraction of trials (Larkman et al., 1991; Allen and Stevens, 1994). The final stimulation amplitude was set to this level (typically 5-16 μA). We only recorded synaptic connections that showed little or no variability in the latency of evoked EPSPs from trial to trial. We then performed 20 ms paired-pulse stimulation at a low frequency (0.2 Hz) for at least 30 sweeps. After recordings, we carefully assessed each sweep by eye in pClamp 9 (Molecular Devices) and included only those sweeps in the final dataset for which an EPSP was evoked following both extracellular stimulation pulses and whose evoked EPSPs were not contaminated by spontaneously occurring EPSPs. As an additional control to ensure that we were stimulating single axons of passage (Allen and Stevens, 1994) and that the synaptic connection remained stable throughout the recording period, we only included synaptic connections when the EPSPs at the end the recording had the identical average amplitude, latency, and shape compared to the first evoked minimal stimulation EPSPs. Our final dataset contained on average 11.2 ± 5 sweeps per synaptic connection (range of 6 to 35 sweeps).

Following the minimal stimulation protocol, we carefully moved the extracellular stimulation electrode to other locations in the L2/3 neuropil to identify different axon fibers forming synapses with the same recorded neuron. Great care was taken not to record from the same stimulation location multiple times, and synaptic connections were only included when their location of stimulation was > 50 µm away from all previous stimulation locations, as assessed in 10x overview images during recordings. At the end of each experiment, we injected current steps into each neuron to characterize its firing pattern as regular-spiking (i.e., putatively excitatory/ pyramidal neuron) or fast-spiking (putatively inhibitory/ interneuron).

### Histology

After recordings, slices were immediately fixed in 15% picric acid, 4% paraformaldehyde, and 0.5% glutaraldehyde in 0.1 M phosphate buffer (PB) overnight. Fixed slices were then washed in PB, incubated in an ascending sucrose ladder for cryoprotection, quickly frozen in liquid nitrogen, and treated in 3% hydrogen peroxide and 10% methanol in phosphate-buffered saline (PBS) to quench endogenous peroxidases. After washing in PBS and tris-buffered saline (TBS), the slices were treated with the Vectastain ABC Kit (Vector Laboratories, catalog # PK-6100, RRID: AB_2336819) in TBS at 4 °C overnight. Following washing in TBS, biocytin was visualized using nickel-diaminobenzidine (Ni-DAB) tetrahydrochloride and hydrogen peroxide treatment, followed by a series of washes in PB to terminate the reaction. Sections were then embedded in Mowiol (Sigma Aldrich) and cover-slipped. Z-stacks of the recovered neurons were imaged under an Olympus BX61 microscope to cross-check the previously determined electrophysiological cell type with anatomy. Pyramidal cells and interneurons were identified on the basis of their dendrite morphology (e.g., spiny dendrites versus smooth dendrites, respectively) and corresponded with the previously recorded regular-spiking firing pattern and fast-spiking firing patterns, respectively.

### Analysis of electrophysiological data

We analyzed each postsynaptic potential evoked with paired-pulse stimulation with Stimfit individually (Guzman et al., 2014) and measured its peak amplitude, coefficient of variation, onset latency (i.e., the time from the onset of the extracellular stimulation artifact to the onset of the evoked postsynaptic potential) and 10% - 90% rise time. The EPSP was defined as the postsynaptic potential evoked by the first pulse of the paired-pulse paradigm, i.e., before STP took place. The paired-pulse ratio was defined as the peak amplitude of the second evoked postsynaptic potential divided by the peak amplitude of the first evoked postsynaptic potential (i.e., the EPSP). Further statistical analyses were done in Matlab (MathWorks) and Prism (GraphPad).

To obtain an unbiased population distribution for a given experiment, we excluded all afferent synaptic connections formed with the postsynaptic neuron in that experiment, but otherwise included all other connections recorded in regular spiking neurons. The cell distribution for a given experiment included all afferent synaptic connections formed with the postsynaptic neuron in that experiment.

We conducted a post-hoc Monte-Carlo power analysis to estimate which effect sizes (i.e., systematic differences between mean EPSP amplitudes or mean paired-pulse ratios between the cell distribution and the population distribution) were detectable given the sample sizes in our dataset. We did this for each experiment individually by bootstrapping new cell distributions with systematically different means and then performing Kolmogorov-Smirnov tests against the population distribution.

Specifically, for the power analysis for paired-pulse ratios, we first formalized the paired-pulse ratio population distribution for each experiment as a normal distribution with the same mean and standard deviation as the experimentally observed paired-pulse ratio population distribution for that experiment. To test which effect sizes were detectable, we then formalized a range of possible underlying generator distributions for the paired-pulse ratio cell distribution for that experiment by varying the mean of the population distribution in steps of ± 0.1 units. By doing so, we designed a range of generator distribution for the paired-pulse ratio cell distribution with systematically different means. For each one of these cell generator distributions, we then drew the same number of random samples that were present in the experimentally observed cell distribution (i.e., between 5 and 8) and ran a Kolmogorov-Smirnov tests against a random sample drawn from the formalized population distribution (containing the same number of entries as the population distribution for that experiment). This analysis was repeated 10,000 times for each cell generator distribution and the statistical power for detecting an effect of a certain size (i.e., the systematic difference in the means between the underlying cell generator distribution and the population distribution) was defined as the fraction of trials that yielded a significant p-value (α = 0.05), see *Results*. The power analysis for EPSP amplitudes was done in an analogous fashion with the only exception that lognormal distributions were used instead of normal distributions, in accordance with our results.

Because our dataset contained 8 experiments for which at least 5 afferent connections were mapped, there were 8 chances for detecting a significant difference between a cell and the population distribution across our experimental series. Thus, a simple binomial model can be used to ask: which systematic difference in paired-pulse ratios should have been observed in at least one of these 8 experiments at the 95% significance levelã To answer this, we computed the probability density functions for obtaining zero as a realization (i.e., the likelihood of observing no significant difference across any of the 8 experiments) of simple binomial functions with N = 8 (i.e., the number of our independent experiments) and P = the average probability of observing a given effect size in a single experiment (as derived above, see *Results*). We then repeated these analyses in an analogous fashion for the EPSP amplitude distributions.

### Conductance-based model of L2/3 neuron

We generated a simplified two-compartment, conductance-based model (Pinsky and Rinsky, 1994; Mainen and Sejnowski, 1996; Larkum, 2004; Yi et al., 2017) of a L2/3 pyramidal neuron in the NEURON software (Hines and Carnevale, 1997). The model neuron consisted of an active soma (diameter of 20 µm) with a Hodgkin-Huxley spiking mechanism and a passive dendrite receiving all synaptic inputs (diameter of 2 µm; length of 100 µm). We set up the model in accordance with experimentally measured passive electrical properties of barrel cortex pyramidal cells, previous models of L2/3 neurons, and our own experimental data.

Because the exact ion-channel compositions for L2/3 neurons are not well established, passive biophysical parameters are routinely modelled as being homogenously distributed in models of L2/3 neurons (Branco et al., 2010; Smith et al., 2013; Ferrarese et al., 2018). It has been determined experimentally that the specific axial resistance (*R*_i_) of pyramidal neurons ranges between 70 Ohm cm to 100 Ohm cm (Stuart and Spruston, 1998), we set *R*i of the dendrite to 100 Ω cm (Wang et al., 2010) to account for the shorter dendrite length and R_i_ of the soma to 1 Ω cm (Wang et al., 2010). In accordance with previous models, we set the specific membrane capacitance (C_m_) of the dendrite to 1.3 μF cm^−2^ to account for dendritic spines, which were not modeled explicitly, and to 1.7 µF cm^−2^ for the soma (Wang et al., 2010). The passive membrane resistivity (R_m_) was set to 8000 Ohm cm^2^ (Branco et al., 2010; Branco and Häusser, 2011; Smith et al., 2013; Ujfalussy et al., 2018), corresponding to a dendritic leak conductance (g_leak_) of 0.126 mS/cm^2^; g_leak_ of the soma was set to 0.0379 mS/cm^2^ (Lajeunesse et al., 2013), and V_m_ was set to -70 mV in accordance with our electrophysiological recordings. To generate action potentials at the soma, we inserted NEURON’s custom Hodgkin-and-Huxley-spiking mechanism at the somatic compartment and used its default values for the active voltage-gated potassium (g^k^ of 0.036 S / cm^2^) and sodium conductance (g^Na^ of 0.12 S / cm^2^).

We inserted 270 synaptic conductances on the dendritic compartment (equidistant to the soma) whose spike times, synaptic weights and short-term plasticity parameters were set as described in the following sections. Briefly, we first constructed 270 spike trains whose pairwise correlation coefficients and firing rates reproduced *in vivo* observations from rodent L2/3 (see *Results*). We then assigned these spike trains with EPSP amplitudes and corresponding paired-pulse ratios that reproduced our *in vitro* data. Synaptic strength was then tuned such that the EPSP amplitudes at the soma of the model neuron matched exactly the somatic EPSP amplitudes we had measured *in vitro* (see below). Importantly, because of this, EPSP amplitudes were independent of the choice of passive biophysical model parameters and there was no need to test the robustness of our simulations towards different sets of passive biophysical model parameters.

### Generating input spike trains with temporal correlations following *in vivo* data

We generated 270 input spike trains whose pairwise correlation coefficients matched the *in vivo* data reported by Cossell et al. (2014), i.e., the minority of (strong) input spike trains exhibited high pairwise correlation coefficients, while the remaining majority of (weak) input spike trains were subsequently less correlated. We first generated a template spike train of 10 s duration that exhibited a sparse and irregular temporal structure by using an inhomogeneous Poisson renewal process and sampling inter-spike interval durations from a gamma distribution (shape k = 1.1, inter-spike interval mean of 40 ms) at 1 ms time steps, which resulted in an average firing rate of 25 Hz. We convolved the template spike train with Gaussian envelopes of different standard deviations (σ_Gaussian_) to generate a set of 270 new spike trains with precisely defined correlation statistics (Azouz, 2005). We divided the 270 inputs into strong (n = 35, i.e., 13 % of inputs) and weak inputs (n = 235, i.e., 87 % of inputs) based on the relationship between EPSP amplitude and short-term plasticity we had found *in vitro* (i.e., synapses with EPSP amplitudes > 2 mV (10 / 74 synaptic connections, i.e., 13.5 %) were exclusively depressing, while synapses with EPSP amplitudes < 2 mV displayed the full range of short-term plasticity). In order to set up these two populations of input spike trains with corresponding temporal correlation statistics, we sampled σ_Gaussian_ from two uniform distributions for strong (σ_Gaussian_ between 5 and 10 ms, n = 35) and weak synaptic inputs (σ_Gaussian_ between 10 and 100 ms, n = 235) (Azouz, 2005). The resulting 270 σ_Gaussian_ values were ranked and assigned to the 270 input spike trains. For each one of the 270 input spike trains, we convolved the spike times of the template spike train with a Gaussian envelope whose standard deviation was set by each spike train’s respective σ_Gaussian_. By doing so, for each spike train, we obtained a 10 s time course consisting of a sum of Gaussian distributions representing the respective spike probability over time. Because of the iteratively increasing σ_Gaussian_, this spike probability distribution for spike trains with increasing indices continuously broadens and flattens with respect to the template spike train. We then generated the discrete spike times for each input spike train by drawing spike times from these time-dependent spike probability distributions using an inhomogeneous Poisson process. The resulting 270 spike trains had continuously lower pairwise correlation coefficients with the template spike train.

Finally, we accounted for the fact that, in barrel cortex *in vivo*, correlated synaptic inputs tend to fire at higher frequencies, while uncorrelated inputs fire at lower rates (O’Connor et al., 2010; Cossell et al., 2015). We parametrized the lognormal firing rate distribution measured by O’Conner et al. (2010) in mouse barrel cortex L2/3 *in vivo* (mean ± s.d.: 4.16 ± 8.33 Hz) and drew 270 random ‘target firing rates’ from it. These values were ranked and assigned to the 270 input spike trains, such that spike trains with higher pairwise correlations with the template spike train also displayed higher target firing rates. We then removed stochastically individual spikes from each input spike train such that the average firing rate of each spike train matched the respective target firing rate.

After the 270 input spike trains had been generated in this manner, we verified that their pairwise correlation coefficients (Cossell et al., 2015) and firing rates (O’Connor et al., 2010) matched experimental data obtained in rodent L2/3 *in vivo* (see *Results*, Fig. 4). This process was repeated 100 times to generate 100 different sets of spike trains to be run in the model.

### Generating EPSP amplitude and corresponding paired-pulse ratio distributions

To assign realistic EPSP amplitudes to the 270 model inputs, we parametrized the EPSP amplitude distribution we measured in regular-spiking neurons *in vitro* with a lognormal distribution (Fig. 1 D, see *Results*) and randomly drew 270 EPSP amplitude values from it. We then generated corresponding paired-pulse ratios for these 270 EPSP amplitudes by parametrized the relationship between the second pulse (EPSP_2_) and the first pulse (i.e., the EPSP amplitude) of the paired-pulse stimulation paradigm that we had recorded *in vitro* (Fig. 1 C) with an exponential decay function. Critically, the jitter of the experimentally recorded EPSP_2_ values around this fitted curve did not differ significantly from a Gaussian distribution (non-parametric Kolmogorov Smirnov Test, p value of 0.48) with a mean ± s.d. of 1.6 * 10^−9^ ± 0.192. This standard deviation captures the natural variance of the ratio of EPSP_2_ to EPSP_1_ and was subsequently used to generate our modeling data. For each of the selected 270 EPSP amplitudes, we first assigned a corresponding EPSP_2_ by using the value predicted by the fitted exponential decay function for the given EPSP_1_ (i.e., the EPSP amplitude). We then added variance to the selected value as a number drawn from a random Gaussian process with a mean of 0 and a standard deviation of 0.192. Finally, we verified that the resulting EPSP distribution, paired-pulse ratio distribution, and their mapping corresponded to our *in vitro* recording data (see *Results*).

### Modeling short-term plasticity dynamically during presynaptic spike trains

The paired-pulse ratio captures a synapse’s short-term plasticity response for two subsequent release events at a stereotypical time interval. To model short-term plasticity dynamically for ongoing activation during spike trains with variable inter-spike intervals, we formalized the short-term plasticity properties of our synapses into a general form by utilizing the widely-used extended Tsodyks-Markram model (Markram et al., 1998; Tsodyks et al., 1998):

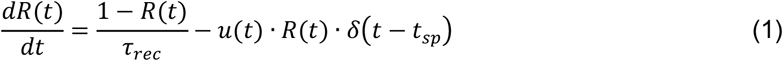

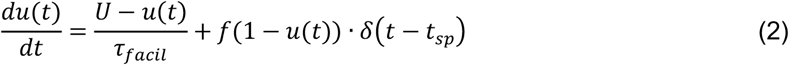

Briefly, short-term depression (equation 1) is modeled as the depletion of the synaptic vesicle pool available for release *R(t)*, with *u(t)* · *R(t)* following a preceding release event at time *t*_*sp*_, which is counterbalanced by vesicle pool recovery at a time constant *τ*_*rec*_. Short-term facilitation (equation 2) is modeled as an increase in release probability *u(t)*, with *f*(1 − *u(t)*) following a preceding spike at *t*_*sp*_, which decays to the baseline release probability *U* with a time constant *τ*_*facil*_. Thus, a continuum of synaptic depression to facilitation can be modeled by specifying the values of the parameter set *Θ =* {*τ*_*rec*_, *τ*_*facil*_, *U, f*} (Costa et al., 2013; Ghanbari et al., 2017).

To do so, we derived *Θ f*or each one of the 270 model synapses as a function of their paired-pulse ratio, as follows. A computationally optimized form of equations (1) and (2) was derived by Costa et al. (2013) by integrating between spikes *n* and *n + 1* at time *Δtn*_*n*_ apart:

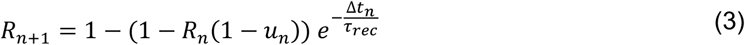

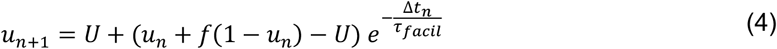

The EPSP amplitude at spike *n*can be calculated as:

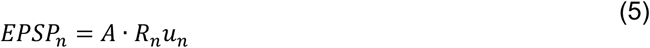

(Markram et al., 1998), where *A* is an adjustable weight parameter that convolves phenomenologically several physiological strength parameters, such as the number of release sites, quantal size, and cable filtering properties. The paired pulse ratio *PPR* is the ratio of the EPSP at spike *n* + 1 and the EPSP at spike *n*:

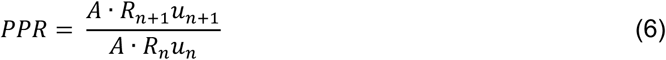

At time *t = 0*, when no preceding spike occurred, the steady-state value of *R*_*n*_ = 1 and of *u*_*n*_ *= U*, and equation (6) can be simplified to:

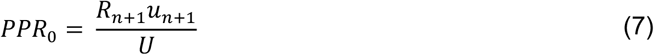

By inserting equations (3) and (4) for *R*_*n*+1,_ *a*nd *u*_*n*+1,_, we can rewrite equation (7) as

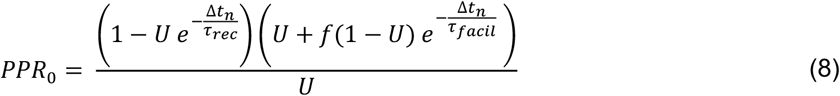

Critically, *PPR*_0_ in equation (8) at *Δtn*_*n*_ = 20 *ms* (i.e., 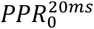) describes exactly our experimental paired-pulse stimulation protocol. This allowed us to obtain a parameter set *Θ f*or each synapse as a function of its 20 ms paired-pulse ratio.

### Defining the short-term plasticity parameter set Θ for each synapse

To do so, we varied Θ on a continuum ranging from strong depression to strong facilitation according to Costa et al. (2013) (Table 1), which resulted in a large dataset of uniquely defined Θs and corresponding 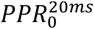 values. For each of our 270 model synapses, we then chose the parameter set Θ, whose resulting 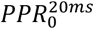 value matched most closely the paired-pulse ratio we had previously assigned to that synapse (see above). By obtaining a unique parameter set Θ for each synapse, we could then compute its *R*_*n*+1,_ and *u*_*n*+1,_ during continuous spike trains using equations (3) and (4), respectively.

**Table 1.** Parameter sets Θ for strongly depressing and strongly facilitation synapses, adopted from Costa et al. (2013).

| Synaptic short-term plasticity | $\tau_{rec}$ | $\tau_{facil}$ | U | f | $PPR_0^{20ms}$ |
| --- | --- | --- | --- | --- | --- |
| Strong depression | 1700ms | 20ms | 0.7 | 0.05 | 0.3 |
| Strong facilitation | 20ms | 1700ms | 0.1 | 0.11 | 1.8 |

### Modeling EPSP amplitude and paired-pulse ratio in the NEURON simulation

We modeled the input synapses by using the *ExpSyn* point process in NEURON, which allows for the synaptic strength to be set precisely by means of a weight parameter. We defined the weight parameter as the product of the desired somatic EPSP amplitude and a scaling factor. To determine this scaling factor, we generated a single test spike for each of the 270 EPSP amplitudes (using its respective desired EPSP amplitude) and measured the resulting somatic EPSP amplitude. We found that the ratio of the desired EPSP / test EPSP was a constant factor across all 270 input synapses, which allowed us to use this ratio as the universal scaling factor.

By deriving paired-pulse ratios using equation (6), we were able to adjust the EPSP amplitudes dynamically in the simulation to incorporate short-term plasticity. To cross-check again that the simulated EPSP amplitudes and short-term plasticity properties reproduced the desired values, each synapse in the NEURON simulation was activated with two pulses at a 20 ms inter-spike interval and the somatic EPSP amplitudes and paired-pulse ratios measured at the soma of the model neuron. Reassuringly, we found that the resulting somatic EPSP amplitude distribution and paired-pulse ratio distribution exactly matched the target distributions we had generated (as described above).

### Modeling the interplay of synaptic strength, short-term plasticity, and temporal correlation in presynaptic spike trains

After the model was set up in this manner, we simulated the somatic voltage response of the model neuron following activation of the 270 input synapses with the corresponding presynaptic spike trains. We convolved the discrete spike times of the output spike train of the model neuron and each one of the 270 input spike trains into continuous functions with an exponential filter (τ = 10ms) (van Rossum, 2001) and computed the pairwise Pearson’s correlation coefficients between each input spike train and the output spike train. Additionally, we quantified the input-output relationship of the model neuron as the probability of spike generation as a function of the number of coincident inputs in the 20 ms time window preceding the output spike. As described in the *Results*, we then manipulated the respective population of active synapses and their synaptic parameters in the simulation to investigate the interplay of synaptic strength, short-term plasticity, and temporal correlation in presynaptic spike trains. We computed mean correlation coefficients, input-output curves and corresponding 95 % confidence intervals by repeating each simulation setup for the 100 sets of spike trains (see above).

## Acknowledgements

We would like to thank Kevan A.C. Martin for his inspiration, support, comments on the manuscript, and funding. We would like to thank Qendrasa Parduzzi for help with developing the NEURON model. As members of the Institute of Neuroinformatics, the authors are signatories of the Basel Declaration. This work was supported by funding from the University of Zurich to Kevan A.C. Martin and by funding of the Swiss National Science Foundation to Gregor Schuhknecht.

## Author Contributions

B.E. and G.F.P.S. designed research,

M.O.B. performed electrophysiology experiments and histology,

M.O.B. and G.F.P.S. analyzed electrophysiology data,

A.G. and B.E. developed the NEURON model,

A.G. and G.F.P.S. analyzed modeling data,

B.E. and G.F.P.S. supervised the work,

G.F.P.S. wrote the paper with input from all authors.

